# Defining developmental diversification of diencephalon neurons through single-cell gene expression profiling

**DOI:** 10.1101/481317

**Authors:** Qiuxia Guo, James Y. H. Li

## Abstract

The embryonic diencephalon gives rise to diverse neuronal cell types, which form complex integration centers and intricate relay stations of the vertebrate forebrain. Prior anecdotal gene expression studies suggest several developmental compartments within the developing diencephalon. In the current study, we utilized single-cell RNA sequencing to profile transcriptomes of dissociated cells from the diencephalon of E12.5 mouse embryos. Through analysis of unbiased transcriptional data, we identified the divergence of different progenitors, intermediate progenitors, and emerging neuronal cell types. After mapping the identified cell groups to their spatial origins, we were able to characterize the molecular features across different cell types and cell states, arising from various diencephalic compartments. Furthermore, we reconstructed the developmental trajectory of different cell lineages within the diencephalon. This allowed the identification of the genetic cascades and gene regulatory networks underlying the progression of the cell cycle, neurogenesis, and cellular diversification. The analysis provides new insights into the molecular mechanism underlying the specification and amplification of thalamic progenitor cells. In addition, the single-cell-resolved trajectories not only confirm a close relationship between the rostral thalamus and prethalamus, but also uncover an unexpected close relationship between the caudal thalamus, epithalamus and rostral pretectum. Our data provide a useful resource for the systematic study of cell heterogeneity and differentiation kinetics within the developing diencephalon.

## INTRODUCTION

Residing in the posterior part of the forebrain, the diencephalon produces distinct classes of neurons, which assemble into multiple nuclei. These functional units are important for linking the anterior forebrain with the rest of the nervous system. For example, the thalamus and epithalamus, which arise from the center of the diencephalon, serve important functions such as learning, motor control, regulation of the sleep-wake cycle, and modulation of emotion and motivation (Hikosaka, 2010; Sherman, 2007). Therefore, understanding the development of the diencephalon is necessary to improve our comprehension of this integral component of the forebrain circuitry.

Based on gene expression patterns and morphological landmarks, the embryonic diencephalon is divided into at least three transverse segments, called the prosomeres (Puelles and Rubenstein, 2003; Rubenstein et al., 1994). During embryogenesis, proliferating progenitor cells in the alar plate of the respective prosomere (p) give rise to the pretectum (p1), the thalamus and epithalamus (p2), and the prethalamus (p3). Additional intra-prosomere subdivisions have been proposed (Ferran et al., 2007; Kiecker and Lumsden, 2005). For instance, the zona limitans intrathalamica (ZLI), which is wedged in between p2 and p3, forms a compartment that is delineated by rostral and caudal cell-lineage restriction boundaries (Zeltser et al., 2001). Importantly, the ZLI acts as an organizer by producing the secreted protein, Shh, to regulate regionalization and patterning of the diencephalon (Kiecker and Lumsden, 2004). Other signals emanating from the roof plate, including those in the Fgf, Bmp and Wnt families, act in concert with Shh to regulate the developmental program within the diencephalon (Chatterjee and Li, 2012; Scholpp and Lumsden, 2010). Despite the progress, our understanding of the pattern formation and neuronal differentiation of the diencephalon remains incomplete. For instance, although much is known about the molecules that define populations of postmitotic neurons arising from different areas of the diencephalon, considerably less is clear about the molecular feature that delineate their respective progenitor populations. Prior studies have revealed highly variable progression in the cell proliferation and neurogenesis, the so-called morphogenetic gradients, across the developing diencephalon (Jones, 2007). Because of the asynchronous development, systematic expression profiling using bulk tissue will likely lack the resolution necessary to gain complete insight into the transcription regulation that underlies the formation of different diencephalic lineages. On the other hand, single-cell gene expression analysis has previously been performed in diencephalon-derived neurons from postnatal mice (Kalish et al., 2018; Phillips et al., 2018; Rosenberg et al., 2018; Zeisel et al., 2018).

In this study, we sought to systematically define the heterogeneity of various cell types and cell states, and to determine the transcriptional landscapes during cell specification and differentiation in the embryonic diencephalon. We performed single-cell RNA sequencing (scRNAseq) of dissociated cells from the mouse diencephalon at embryonic day (E) 12.5. Our unbiased scRNAseq allowed characterization of the molecular features across different cell types and cell states that arise from various diencephalic compartments. Furthermore, we inferred developmental trajectories of diverse cell lineages within the diencephalon. This led to identification of the genetic cascades and gene regulatory networks underlying the progression of the cell cycle, neurogenesis, and cellular diversification. To the best of our knowledge, this is the first reported scRNAseq analysis of mouse embryonic diencephalon. Our data provide a valuable resource for systematic study of cell heterogeneity and differentiation kinetics within the diencephalon.

## RESULTS AND DISCUSSION

### Classification of cell groups of the mouse diencephalon

We used a mouse line, in which bicistronic *creER-ires-EGFP* transcripts are expressed from the G*bx2* locus so that thalamic neurons specifically produce both creER and enhanced green fluorescence protein (EGFP) (Chen et al., 2009). Utilizing EGFP as a guide, we dissected the thalamus and surrounding tissues from mouse embryos at E12.5. Using the Chromium Drop-Seq platform (10x Genomics), we profiled the transcriptome for over 7,500 single cells. After applying quality filters, we obtained a dataset with 7,365 cells and 14,387 genes for subsequent analysis. Using the Seurat algorithm (Butler et al., 2018; Satija et al., 2015), we partitioned the 7,365 cells into 18 clusters, which were visualized with t-distributed stochastic neighbor embedding (t-SNE; Fig. 1A). Differential gene expression analysis identified genes that were significantly enriched in each cell cluster (Fig. 1B and Supplementary Table S1). We used a set of cell-cycle related genes (Tirosh et al., 2016) to calculate cell-cycle scores and thereby to assign cell-cycle status (G2/M, S, or postmitotic) to each cell (Fig. 1C). In t-SNE projections, the distribution of cells with various mitotic statuses showed a trend reflecting the progression from proliferating progenitors to postmitotic cells (Fig. 1C). Inspection of the average gene counts revealed a trend of decreasing transcript levels from progenitors to postmitotic cells (Fig. 1D), indicating that the dividing progenitors have higher gene counts than their progeny. We classified cluster 11 as low-quality cells, as they had much lower gene counts than the others, and contained few cluster-specific genes (Fig. 1B and D). Besides brain cells, we recovered non-neural cell types, such as endothelial cells (cluster 17) and microglia (cluster 18). Hierarchical analysis classified the 18 cell clusters into five main groups: postmitotic neurons, neuronal precursors or committed progenitors, neural progenitors, non-neural cells, and low-quality cells (Fig. 1E). We identified the markers that were common for progenitors (*Id3* and *Ptn*), newly committed progenitors (*Gadd45g* and *Hes6*), and postmitotic neurons (*Stmn2* and *Gap43*; Fig. 1F and Table S1). Therefore, our scRNAseq data have quantified the heterogeneity of cells within the mouse diencephalon at E12.5.

**Figure 1.**
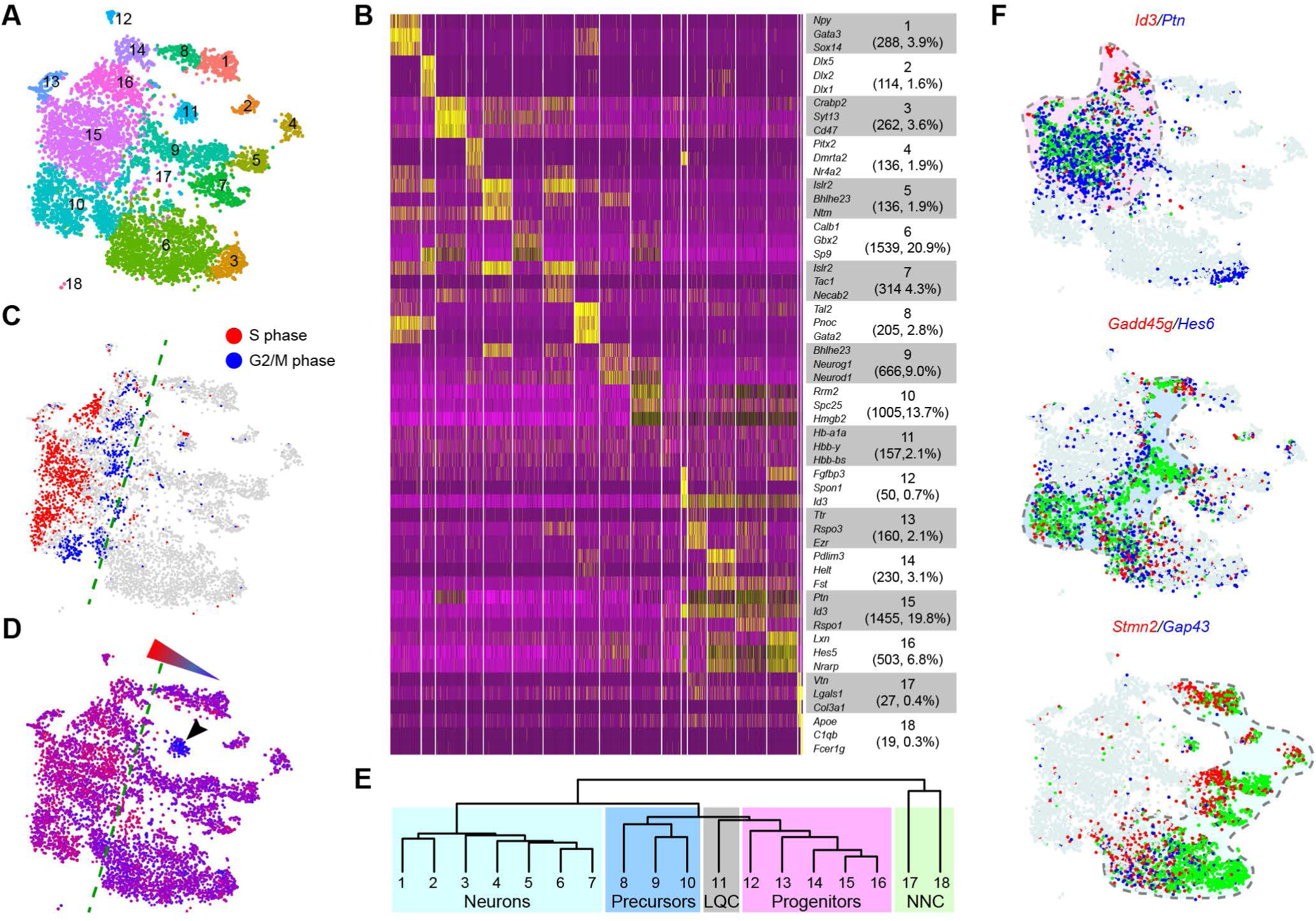
Identification of major cell groups in E12.5 mouse diencephalon by scRNAseq. (**A**) Visualization of 18 classes of cells using t-distributed stochastic neighbor embedding (t-SNE). Each dot represents a single cell, similar cells are grouped and shown in color. (**B**) Heatmap showing expression of marker genes across cell groups. The number and percentage of cells are shown in the bracket under the cluster number. (**C, D**) t-SNE plots showing the inferred cell cycle phase (C) and average gene counts (D). The dashed lines delineate between proliferating cells (to the left) and postmitotic cells (right). In D, the arrowhead indicates the low-quality cells, and the triangle denotes the gradient of gene counts. (**E**) Dendrogram showing the relationship between cell groups recovered by scRNAseq. Abbreviations: LQC, low-quality cells; NNC, non-neural cells. (**F**) Expression of the genes marking cell clusters corresponding to neural progenitors, neuronal precursors, and postmitotic neurons, respectively. ≈

### Characterization of the molecular feature of postmitotic neurons

Next, we related postmitotic cell groups to their endogenous positions by inspecting RNA *in situ* hybridization data in the Allen Developing Mouse Brain Atlas (Thompson et al., 2014). By comparing the expression of at least two markers in t-SNE projections and *in situ* hybridization on serial sagittal sections of mouse embryonic brains, we assigned cell clusters 3 to caudal thalamus, cluster 1 to rostral thalamus, cluster 2 to prethalamus, cluster 4 to ZLI, cluster 7 to epithalamus, and cluster 5 to pretectum (Fig. 2A-F). Clusters 6, 8, and 9 apparently represented intermediate cell states in transition to more differentiated cells of clusters 3, 1, and 5/7, respectively (see below). As the Allen Developing Mouse Brain Atlas has limited information of E12.5 mouse brains, we curated a list of markers for different diencephalic cell types at E12.5 based on published studies (Chatterjee et al., 2012; Delogu et al., 2012; Mallika et al., 2015; Suzuki-Hirano et al., 2011; Virolainen et al., 2012). Examination of these known markers confirmed our annotation of postmitotic cell clusters (Fig. 2G-I).

**Figure 2.**
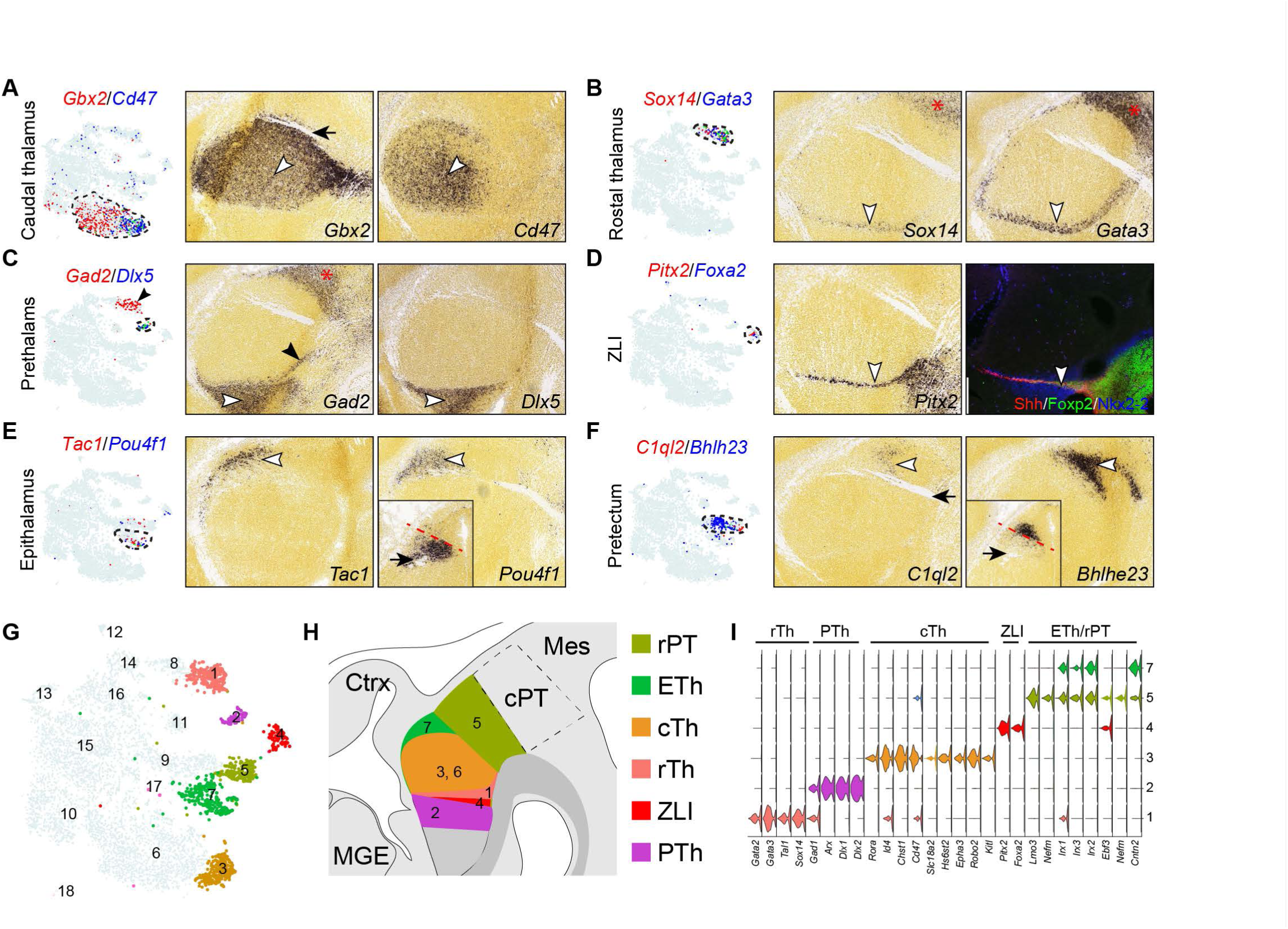
Relating postmitotic cell groups to their spatial positions in the diencephalon. (**A-F**) Expression of cell-specific markers with t-SNE plotting (left) and on sagittal sections of E13.5 mouse diencephalon (right, from Allen Developing Mouse Brain). The white arrowhead indicates the marker expression domain; the black arrow denotes the fasciculus retroflexus; the black arrowheads in C show that Gad2 is expressed in the rostral thalamus, as well as the prethalamus; the asterisks indicate the caudal pretectum, which exhibits a similar gene expression profile as the rostral thalamus. Insets in E and F show the expression of *Pou4f1* and *Bhlhe23* on coronal sections of E13.5 diencephalon; the red dashed line separates their expression domains. (**G**) t-SNE projections of diencephalic neurons. (**H**) Schematic summary of the postmitotic neurons identified by scRNAseq in their endogenous positions in the diencephalon. (**I**) Violin plot showing expression of genes that are known to mark different diencephalic neurons at E12.5. Abbreviations: ETh, epithalamus; cTh, caudal thalamus; PTh, prethalamus; rPT, rostral thalamus; rTh, rostral thalamus; ZLI, zone limitans intrathalamica.

**Figure 3.**
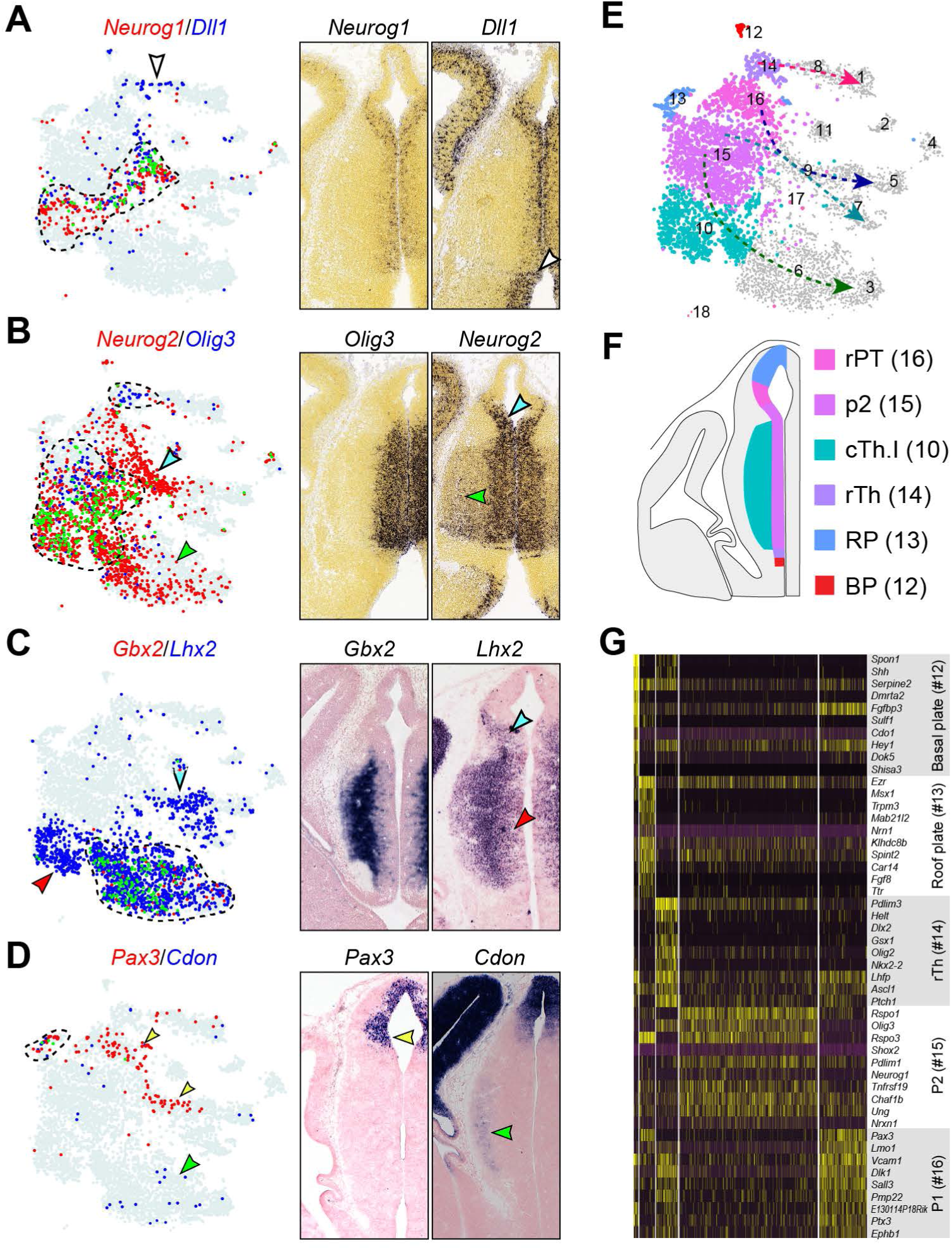
Relating progenitor cell groups to their original positions in the diencephalon. (**A-D**) Expression of cell-specific markers on t-SNE plots (left) and on coronal sections of mouse diencephalon (right) at E13.5 (A and B, from Allen Developing Mouse Brain) and E12.5 (C and D). On t-SNE plots, the cells that express a given marker are shown in red or blue as indicated, while those express both are shown in green and outlined by dashed lines. The cell group that expresses a single marker is indicated by an arrowhead with matching color on both t-SNE plots and sections. (**E**) t-SNE plotting of diencephalic progenitor cell groups. The dashed lines show developmental trajectories. (**F**) Schematic representation of the scRNAseq-identified progenitor groups within the diencephalon. (**G**) Heatmap showing markers for different progenitor cell groups. Abbreviations: BP, basal plate; ETh, epithalamus; cTh, caudal thalamus; p1/2, prosomere 1/2; PTh, prethalamus; RP, roof plate; rPT, rostral thalamus; rTh, rostral thalamus.

### Identification of spatial gene expression patterns in the roof plate and p2

The epithalamus and the thalamus arise from the dorsal and ventral parts of p2 (Chen et al., 2009; Mallika et al., 2015). Interestingly, cell clustering could not reliably subdivide the presumed p2 progenitor cells (cluster 15), suggesting that the epithalamic and thalamic progenitors share a similar molecular feature. Alternatively, these two groups of progenitors might contain gradual expression changes of relatively few genes. To test the latter hypothesis, we explored the “spatial structure” in our scRNAseq data with different analytical tools.

*Fgf8* is expressed in two narrow longitudinal stripes flanking the roof plate overlaying p1-2, whereas *Wnt1* is expressed at the roof plate covering the p1 and mesencephalon (Martinez-Ferre and Martinez, 2009; Martinez-Ferre et al., 2013). In t-SNE projections, the roof-plate cells (cluster 13) was juxtaposed with the p2 cells (cluster 15; Fig. 3E). *Fgf8* and *Wnt1* were detected in two small discrete subgroups within the roof-plate cluster (Fig. S1D and E). Moreover, *Wnt1*^+^ cells were embedded with *Pax3*^+^ cells, indicating a p1 origin of these cells (Fig. S1D). A decreasing gradient of *Rspo3* expression was detected from the *Fgf8*^+^ cells to the rest of p2 progenitor cells, recapitulating the endogenous expression pattern (Fig. S1E and F). These observations suggest that spatial information is preserved among cells in t-SNE projections. We next applied trendsceek, a statistical method, which tests points (cells in low-dimensional projections) in a pairwise manner to identify genes that exhibit distance-dependent expression in single cell gene expression dataset (Edsgärd et al., 2018). Trendsceek identified 92 genes with significant spatial trends (p < 0.01, Benjamini-Hochberg adjusted), including at least six different patterns exemplified by *Ttr*, *Fgf8*, *Bmp4*, *Pax3*, *Irx3*, and *Olig3*, respectively (Fig. 4A-F). Remarkably, cells expressing these genes occupied different spatial regions within the diencephalon, as well as in t-SNE projections (Fig. 4K and L), validating the trendsceek analysis. *Ttr*, which encodes a transporter protein that carries thryroxin and retinol, is normally expressed in precursor cells of choroid plexus (Herbert et al., 1986). Therefore, we classified the *Ttr*^+^ cell group as choroid plexus epithelium.

**Figure 4.**
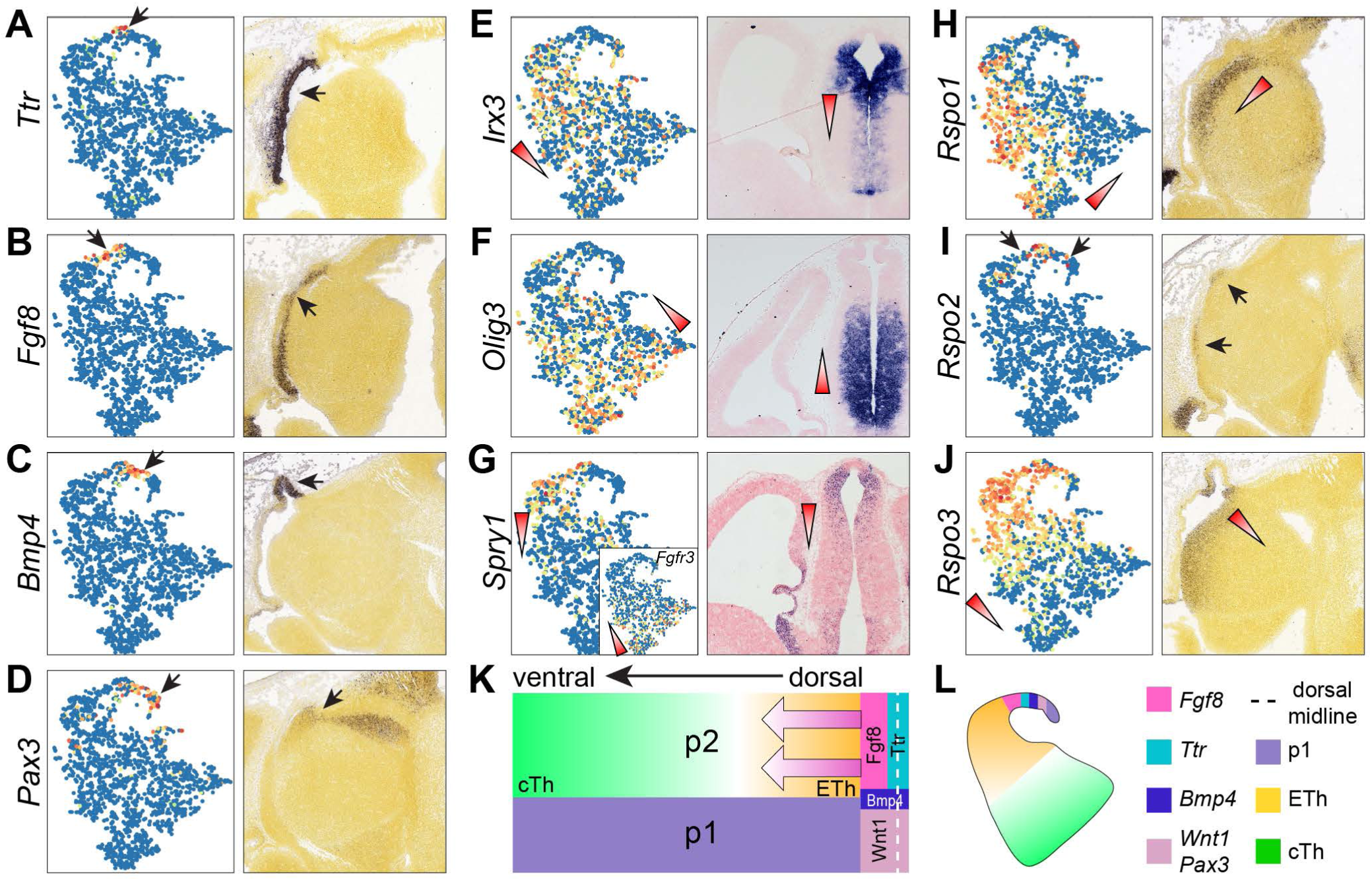
Inference of spatial gene expression patterns with scRNAseq data. (**A-J**) Spatial expression of significant genes identified by trendsceek on scRNAseq data projected with t-SNE (left) and on sagittal brain sections (right). Expression levels are scaled from 0 (blue) to 1 (red); arrows indicate the matching expression domain on the section and cells on the t-SNE; triangles denote the expression gradient. Images in A-G are from Allen Developing Mouse Brain Atlas. (**K** and **L**) Schematic representation of the spatial expression patterns in the diencephalon (K) and t-SNE (L). The arrow indicates the morphogen gradients of Fgf8 and Wnt/ß-catenin signaling.

We uncovered 36 genes that displayed opposing gradients in p2 progenitors along the presumptive dorsoventral axis similar to the expression patterns of *Irx3* and *Olig3* (Fig. S2B and C). Some of these genes have been implicated in the p2 regionalization. For instance, *Irx3* and *Pax6*, which exhibit a dorsal^high^-vental^low^ gradient (Fig. 4E, S2A, and S2B), are known to establish the differential competence for Shh-signaling in the patterning of the p2 progenitor zone (Robertshaw et al., 2013). *Tcf7l2*, which shows a ventral^high^-dorsal^low^ gradient (Fig. S2B), is important for thalamus development as its deletion results in a thalamic-to-epithalamic fate conversion (Lee et al., 2017b). Collectively, our data suggest that the prospective epithalamus and thalamus arise from two p2 progenitor domains with graded, rather than discrete, gene expression differences.

It has been shown that *Fgf8* signaling induces *Spry1* but represses *Fgfr3* (Liu et al., 2003). We detected opposing gradients of *Spry1* (dorsal^high^) and *Fgfr3* (ventral^high^) (Fig. 4G). This indicates the presence of graded *Fgf8* activity in the p2 progenitor domain, in agreement with the known function of *Fgf8* in patterning of p2 (Martinez et al., 1999). A prior report has demonstrated a complex pattern of Wnt/ß-catenin signaling, with lower activity in the ventral and rostral regions, in the p2 domain (Bluske et al., 2009). However, the pattern of Wnt/ß-catenin activity does not correlate with the expression pattern of known Wnt ligands. Trendsceek analysis uncovered significant spatial trends of *Rspo1-3*, which code for secreted proteins that potentiates Wnt/ß-catenin signaling (de Lau et al., 2014). Notably, *Rspo1* is expressed in a caudal^high^-rostral^low^ gradient; *Rspo2* is expressed near the roof plate; *Rspo3* is expressed in a dorsal^high^-ventral^low^ gradient (Fig. 4H-J). Therefore, the differential activity of Wnt/ß-catenin may be governed by the combined function of Rspo1-3 proteins in the p2 progenitor domain. Collectively, our findings suggest that the morphogen gradients of Fgf8 and Wnt/ß-catenin signaling may orchestrate the graded gene expression in p2 progenitor zone, leading to the formation of the epithalamus and thalamus from the dorsal and ventral parts of p2, respectively.

### Progenitor cells in the caudal thalamus display a unique pattern of cell cycling and neurogenesis

Prior studies have demonstrated the existence of a unique and significant population of basal progenitors in the caudal thalamus (Smart, 1972; Wang et al., 2011). As expected, we recovered abundant basal progenitors (cluster 10) presumably from to the caudal thalamus (Fig. 3E). This cluster contained cells in S and G2/M phases, despite the robust expression of the proneural genes *Neurog1* and *Neurog2* (Fig. 1C, 2A, and 2B), indicating that these neurogenic progenitors are highly proliferative. It has been shown that basal progenitors in thalamus depends on *Neurog1/2* and *Pax6* (Wang et al., 2011). However, as these genes are expressed in other regions that lack basal progenitors, other unknown factors must be involved to prevent the cell cycle exit of basal progenitors in the thalamus.

To characterize the progression of cell proliferation and neurogenesis in the caudal thalamus, we digitally isolated cells that presumably originated from this domain (clusters 3, 6, 10 and 15), and applied the Monocle 2 algorithm to determine the developmental continuum according to pseudotemporal order or pseudotime (Qiu et al., 2017). For comparison, we performed the same analysis for the rostral pretectal cells (clusters 5, 7, 9, and 16). Inspection of pseudotime and expression of known markers, *Id3* (apical progenitors), *Neurog1* (basal progenitors), *Gbx2* (newborn neural precursors), and *Hs6st2* (postmitotic neurons), showed that Monocle correctly ordered cells according to the developmental trajectory of the thalamic lineage (Fig. 5A-C). Interestingly, the apical progenitor (*Id3*^+^) and basal progenitor (*Neurog1*^+^) groups each contained one group of M-phase cells, and two separate groups of S-phase cells in t-SNE projections (Fig. 5D), suggesting that both apical and basal progenitors in the caudal thalamus undergo actively symmetric division. Ordering cells according to pseudotime revealed sequential waves of gene expression and temporal coupling between cell cycle and neurogenesis in the caudal thalamus and rostral pretectum (Fig. 5E and F). Remarkably, the expression of the M-phase (*Bir5*) and S-phase (*Atad2*) genes (Tirosh et al., 2016) displayed at least two out-of-phase waves, overlapping with the induction of *Neurog1* and *Neurog2* (Fig. 5E). By contrast, the expression of the cell-cycle genes and apical progenitor markers precipitously decreased before the induction of *Neurog1* and *Neurog2* in the rostral pretectum (Fig. 5F). We detected coexpression of *Insm1* and *Neurog1/2* in both the caudal thalamus and rostral pretectum (Fig. 5E and F), indicating that *Insm1* is not specific for basal progenitors as suggested previously (Wang et al., 2011). However, the induction of *Cdkn1c*, which codes for a cyclin dependent kinase inhibitor (p57^kip2^) important for neuronal cell cycle exit (Dyer and Cepko, 2001; Gui et al., 2007), was delayed in the caudal thalamus compared to that in the rostral pretectum (Fig. 5E and F). Therefore, we demonstrate the unique pattern of the cell cycle progression and neurogenesis in the caudal thalamus, in agreement with previous studies (Smart, 1972; Wang et al., 2011). Importantly, our data suggest that the delayed induction of *Cdkn1c* is involved in the significant expansion of basal progenitors in the caudal thalamus, which may be responsible for the generation of vast number and diversity of thalamic neurons in mammals.

**Figure 5.**
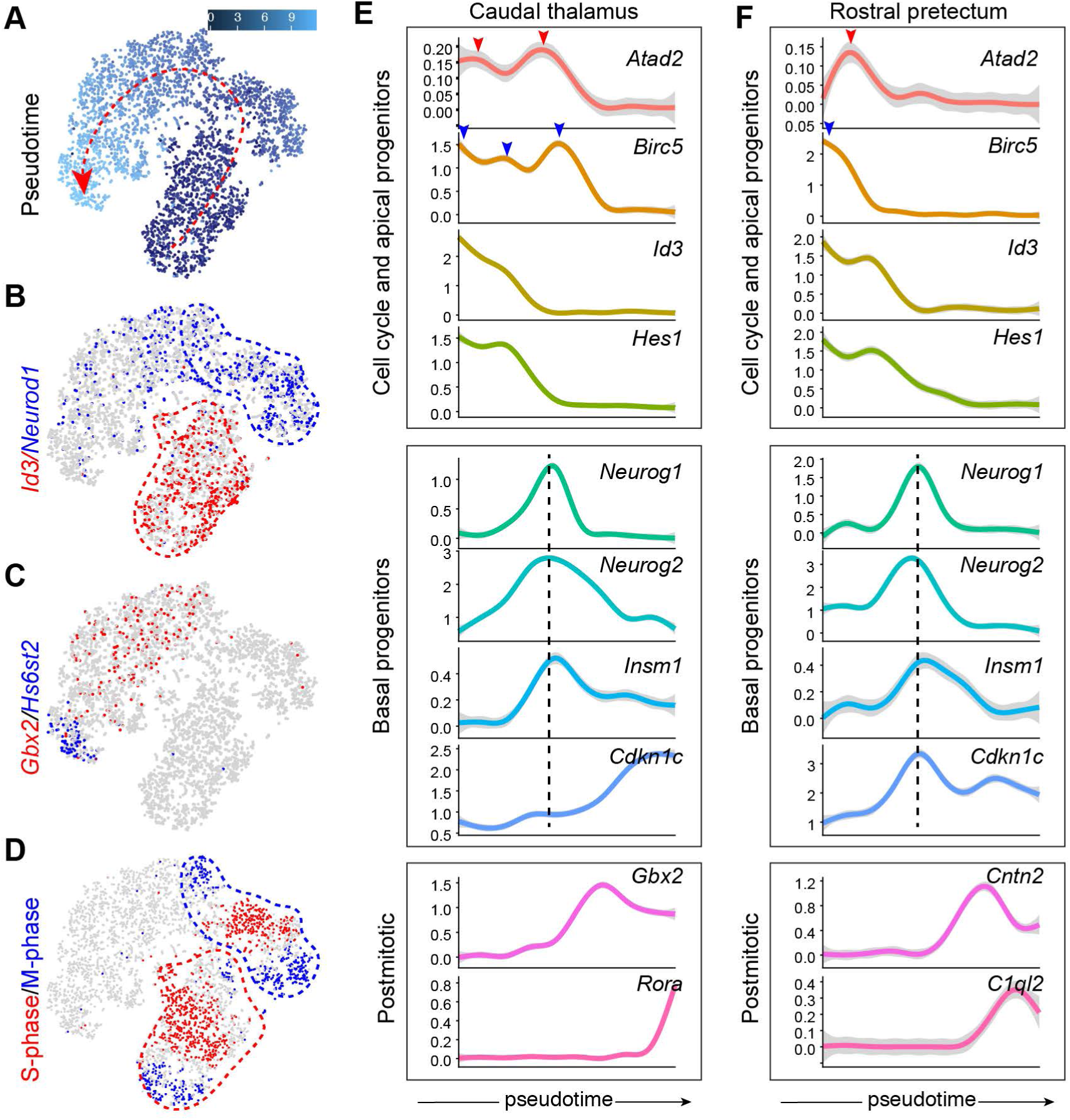
Examination of the unique pattern of cell cycle exit and neurogenesis of thalamic progenitors. (**A-D**) t-SNE plots showing pseudotime (A), gene expression (B and C), and inferred cell cycle states (D) of cells originating from the caudal thalamus. The dashed arrow indicates the increase of pseudotime predicated by Monocle. (**E** and **F**) Temporal expression profiles of the genes that represent cell cycle and apical progenitors, basal progenitors, and postmitotic neurons, respectively, in the caudal thalamus (E) and rostral pretectum (F). The arrowheads indicate the peak expression of genes representing the S (red) and M (blue) phases; the vertical dashed lines show the pseudotime position that corresponds to the peak of *Neurog1/2* expression.

### Inference of developmental trajectories of lineage specification

To reconstruct development trajectories of various diencephalic cell types, we applied the URD algorithm (Farrell et al., 2018) to our scRNAseq data. Building on an extension of the diffusion map framework (Haghverdi et al., 2015; Haghverdi et al., 2016), URD uses discrete random walks and graph searches to order cells in pseudotime and reconstruct complex developmental trajectories (Farrell et al., 2018). We assigned apical progenitors (mostly clusters 15 and 16) as the root, and postmitotic neurons of clusters 1-5 and 7 as tips, resulting a multi-branching trajectory (Fig. 6A). The hierarchical tree was visualized with a force-directed layout based on cells’ visitation frequency by the random walks from each tip (Fig. 6B). Inspection of the expression of known markers showed progressive changes along consecutive segments of the respective trajectory as expected (Fig. 6C and Fig. S3), suggesting that we had successfully reconstructed the developmental trajectory of the diencephalon.

**Figure 6.**
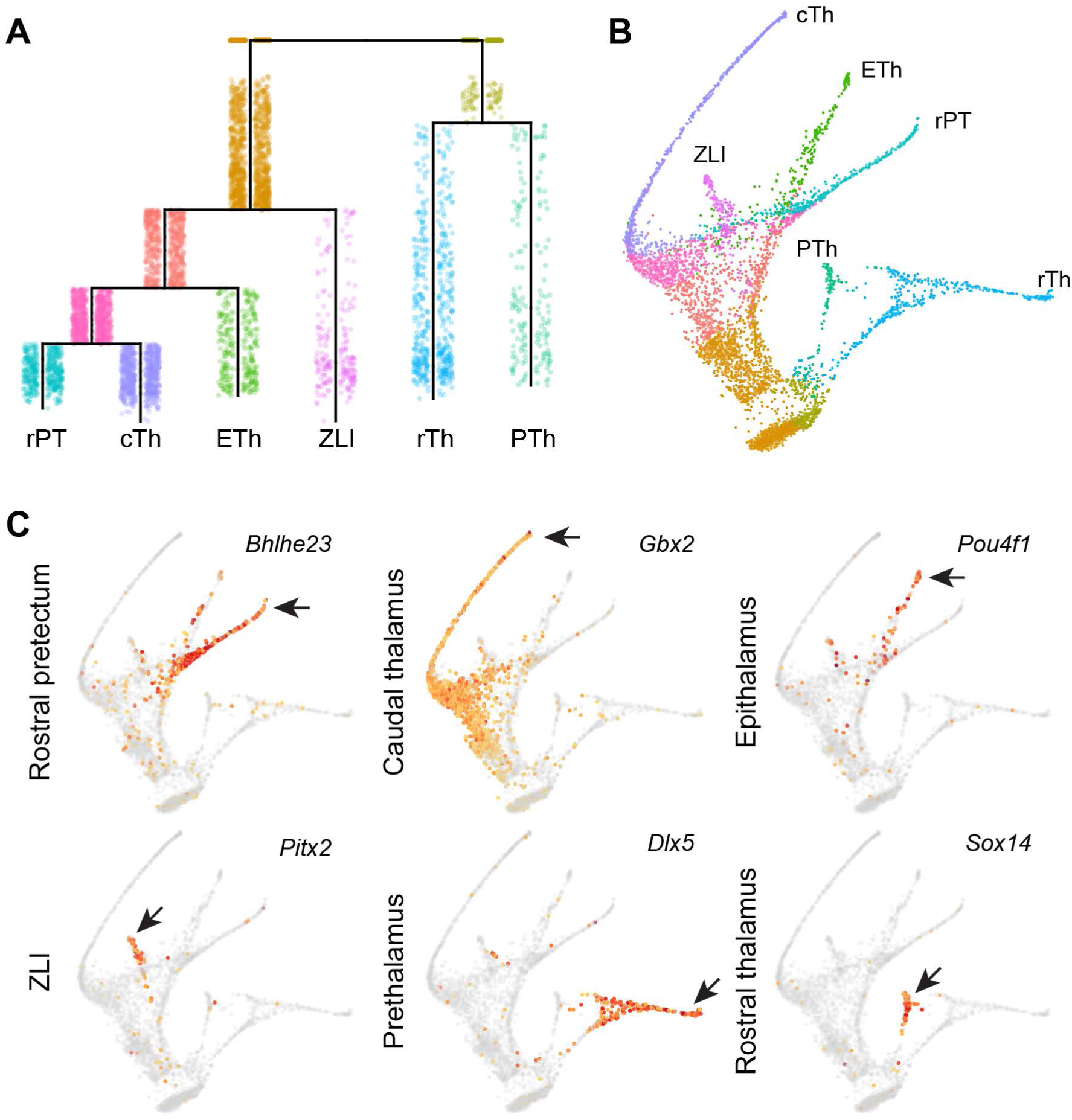
Reconstruction of developmental trajectories of the diencephalon. (**A**) URD-inferred hierarchical tree of diencephalic lineages. (**B**) Force-directed layout of the trajectory tree colored by segments as shown in A. (**C**) Expression of lineage markers overlaid on the trajectory tree. The arrows indicate the robust expression in each specific lineage. Abbreviations: cTh, caudal thalamus; ETh, epithalamus; PTh, prethalamus; rPT, rostral pretectum; rTh, rostral thalamus; ZLI, zona limitans intrathalamica.

The reconstructed tree closely reflects the development of different populations of diencephalic neurons. For example, the URD-inferred tree predicts a close relationship between the rostral thalamus and prethalamus (Fig. 6A and B). This is in agreement with the notion that these two regions may share a common pool of progenitors but give rise to different subtypes of GABAergic neurons (Kataoka and Shimogori, 2008). Indeed, GABAergic precursors that arise from the rostral thalamus or pretectum retain latent plasticity for subtype identity upon cell cycle exit; removal of various transcription factors, such as *Gata2*, *Helt* and *Ascl1*, *Tal2*, and *Dlx1/2* results in cell fate switching between rostral thalamus and prethalamus (Delogu et al., 2012; Lee et al., 2017a; Sellers et al., 2014; Song et al., 2015; Virolainen et al., 2012). Furthermore, the predicted close relationship between the epithalamus and the thalamus is also in line with the fact that they both arise from the p2 domain (Puelles and Rubenstein, 1993; Rubenstein et al., 1994). In addition, we and others have reported that the deletion of *Gbx2* or *Tlc7l2* causes cell-fate conversion from thalamic neurons to those of an epithalamic fate (Lee et al., 2017b; Mallika et al., 2015).

The predicted closed relationship between the rostral pretectum and the thalamus or the epithalamus is somewhat unexpected as they belong to p1 and p2, respectively. However, evidence from genetic studies in mice and fish has recently suggested a latent cell-fate plasticity between the thalamus- and pretectum-derived neurons. For example, the deletion of *lhx2/9* or increased Wnt signaling causes respecification of the thalamic to a pretectal fate in the zebrafish embryo (Peukert et al., 2011). Conversely, ectopic expression of *Shh* can convert the pretectum into a thalamic fate (Kiecker and Lumsden, 2004; Vieira and Martinez, 2006; Vieira et al., 2005). Finally, although it has been demonstrated that the thalamic neurons are converted to an epithalamic fate in mice lacking *Gbx2* or *Tcf7l2* (Lee et al., 2017b; Mallika et al., 2015), our new evidence suggests that the rostral pretectum-specific markers were also ectopically increased the thalamus in the absence of *Gbx2* and *Tcf7l2* (see below). Collectively, these observations support the view that the thalamus and pretectum, as least the rostral part of the pretectum, are closely related.

Therefore, the single-cell-resolved trajectories not only confirm a close relationship between the rostral thalamus and prethalamus, but also uncover an unexpected close relationship between the caudal thalamus, epithalamus and rostral pretectum. It is worth noting that the deduced lineage relationship is based on gene expression. The actual ancestor-progeny relationship can be determined in the future by the emerging technology that combine lineage tracing and scRNAseq (Kester and van Oudenaarden, 2018).

### Determination of gene expression landscapes along developmental trajectories

To determine gene expression dynamics in the molecular specification of diencephalic cell lineages, we searched for genes whose expression levels significantly changed along a given URD-recovered trajectory. We identified a total of 647 such so-called “cascade genes” in each trajectory leading to the formation of the epithalamus (n = 313), caudal thalamus (n = 361), prethalamus (n = 100), rostral pretectum (n = 335), rostral thalamus (n = 94), and ZLI (n = 157; Supplementary Table S2). For the cascade genes in each lineage, we applied impulse response fitting to determine their temporal dynamics (Fig. 7A-F). This analysis also allowed us to identify markers specific for each lineage (Fig. 7A-F and Table S2). Assessing the expression of 20 thalamus-specific markers predicted by URD showed that they correctly highlighted the caudal thalamus branch in the trajectory tree (Fig. S4). Notably, 13 of them (65%) have been shown to be specific for the thalamic neurons according to previous studies (Mallika et al., 2015; Suzuki-Hirano et al., 2011), indicating the accuracy of lineage-specific markers recovered by URD.

**Figure 7.**
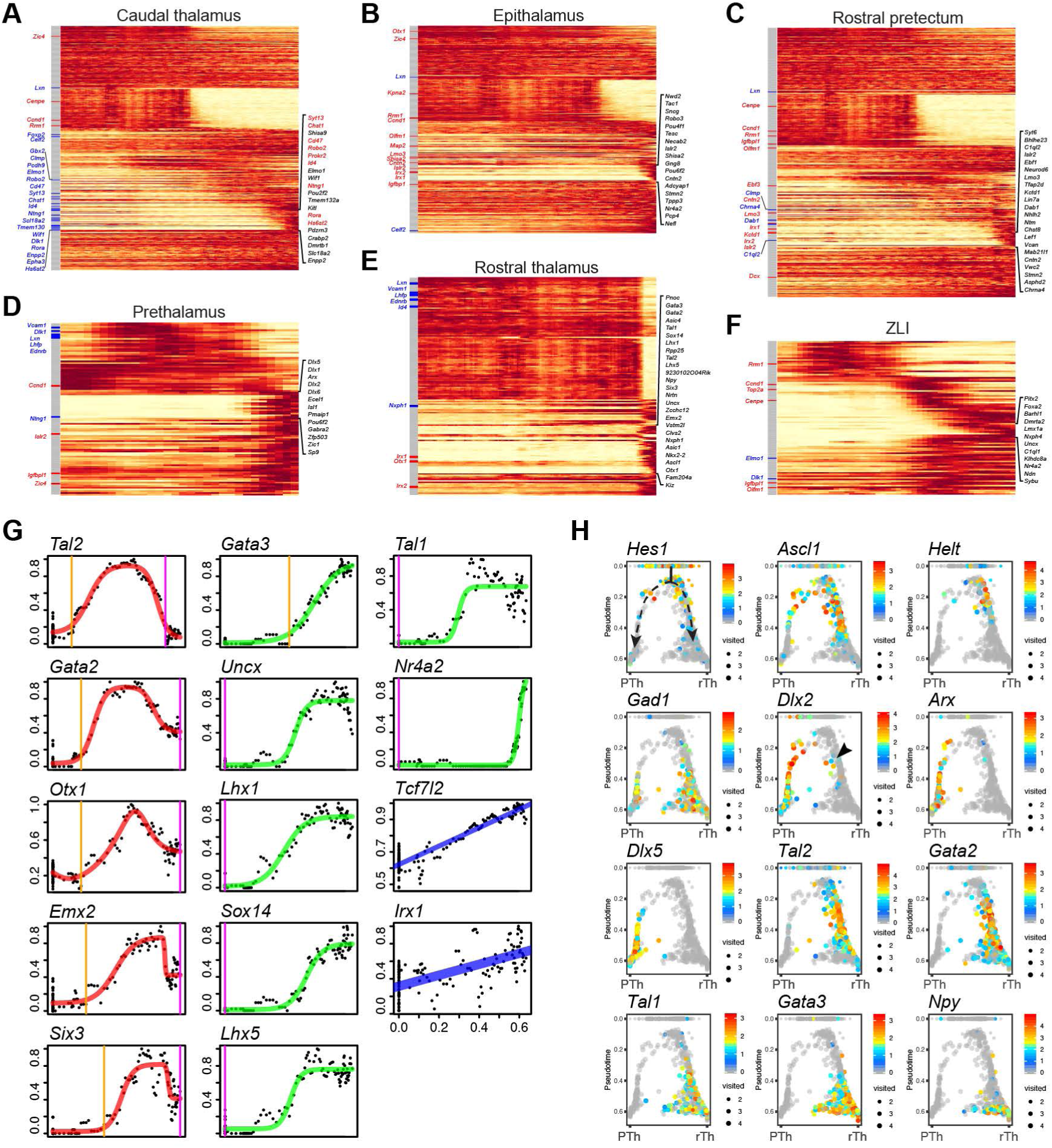
Gene expression cascades involved in the development of different diencephalic lineages. (**A-F**) Heatmap showing gene expression dynamics in different diencephalic lineages. Pseudotime increases from left to right. The lineage-specific genes are shown to the right. To the left are the genes that are significantly altered in the thalamus by *Gbx2* deletion (red, up-regulated; blue down-regulated; grey unchanged). (**G**) Temporal dynamics of 14 transcription factors that exhibit the most significant changes in the specification of rostral thalamic neurons. Each dot represents the moving-window average of expression levels; the line shows impulse response fitting: blue for genes that change at a constant rate, green for a single sigmoid impulse, and red for a convex impulse; vertical lines indicate the onset (orange) and offset (magenta) of gene expression. (**H**) Expression of key developmental regulators overlaid on the branch point forming the rostral thalamus (left) and the prethalamus (right), respectively. The dashed line indicates the lineage bifurcation and trajectory; the arrowhead shows the presence of *Dlx2* expression in rostral thalamic cells near the branch point.

It has been demonstrated that *Gbx2* is essential for the development of the thalamus (Chatterjee et al., 2012; Chen et al., 2009; Miyashita-Lin et al., 1999). Through microarray transcriptome profiling, we have identified a set of genes whose transcription is significantly altered in the mouse thalamus at E12. 5 as a consequence of the deletion of *Gbx2* (Mallika et al., 2015). Remarkably, the down-regulated genes showed significant enrichment for URD-predicted markers of the caudal thalamus (hypergeometric tests, Bonferroni corrected p = 1.37 x 10^-23^), whereas the up-regulated genes were significantly enriched for those of the epithalamus (p = 2.41 x 10^-05^, Fig. 7A, and B). This is in agreement with the conclusion that the loss of *Gbx2* results in an epithalamus-to-thalamic cell fate change (Mallika et al., 2015). Unexpectedly, we found that the up-regulated genes in the *Gbx2*-deficient thalamus were also enriched for rostral pretectum markers (p = 4.51 x 10^-05^; Fig. 7C). Therefore, our data have extended our previous findings by showing that the loss of *Gbx2* causes respecification of the thalamic fate to a cell fate of the rostral pretectum, as well the epithalamus. We observed that many genes, such as *Irx2*, *Islr2*, *Lmo3*, and *Ebf3* that were reported to be expressed in the thalamus lacking *Tcf7l2* (Lee et al., 2017b), were indeed markers for the rostral pretectum (Fig. 7C). This suggests that the loss of *Tcf7l2* may result in a partial cell fate switch from thalamus to rostral pretectum. Collectively, these observations further support the idea that the formation of the epithalamus, caudal thalamus, and rostral pretectum are closely related in transcriptional regulation.

To gain insight into the molecular underpinnings of cell fate decision, we generated impulse models for transcriptional regulators that exhibited the most dynamic changes in each diencephalic lineage (Fig. S5). We identified 14 such transcription factors in association with the development of the rostral thalamus (Fig. 7G). Remarkably, the predicted temporal expression profile of these genes recapitulated their known expression behavior (Delogu et al., 2012; Sellers et al., 2014; Virolainen et al., 2012). Many of these proteins have been demonstrated to play an essential role in GABAergic neuron differentiation in the rostral thalamus or in other brain regions (Achim et al., 2012; Delogu et al., 2012; Lee et al., 2017a; Sellers et al., 2014; Song et al., 2015; Virolainen et al., 2012). Our analyses also revealed dynamic expression of several transcription factors, such as *Otx1*, *Emx2*, *Six3*, *Uncx*, *Nr4a2*, *Tcf7l2* and *Irx1*, without described roles in the rostral thalamus. These genes would be candidates for future genetic studies.

For validation, we focused on the expression of known molecules at the branch point separating the prospective rostral thalamus and prethalamus (Fig. 7H). As expected based on previous studies (Lee et al., 2017a; Sellers et al., 2014; Song et al., 2015; Virolainen et al., 2012), most cells fell along the bifurcation from progenitors into rostral thalamic and prethalamic fates with sequential gene expression changes corresponding to the progression of lineage development (Fig. 7H). Notably, a small fraction of cells that expressed *Dlx2* were found at the early stage of the rostral thalamic lineage, indicating a latent fate plasticity of these cells. Therefore, repression of *Dlx2*, possibly by the *Gata2*, is important for the commitment of the rostral thalamic fate as shown previously (Sellers et al., 2014; Virolainen et al., 2012).

Collectively, our analysis has successfully determined the gene regulation dynamics associated with the development of different diencephalic lineages. Our data provide a valuable resource for interrogation of the cell fate specification and differentiation in the developing diencephalon.

### Examination of transcriptional networks in single-cell-resolved trajectories

Next, we investigated how the URD-inferred developmental trajectories are characterized by gene regulatory networks regulated by transcription factors. We applied the single-cell regulatory network inference and clustering (SCENIC) (Aibar et al., 2017) pipeline to uncover the gene regulatory network that underlies the cell-state transition in different diencephalon lineages. We first identified sets of genes that were coexpressed with transcription factors using the GEINE3 algorithm (Huynh-Thu et al., 2010). We then applied RcisTarget (Aibar et al., 2017) to select the coexpression modules, called regulons, which exhibited significant enrichment for binding motifs of the corresponding transcription factors. This resulted in 230 regulons in our dataset. The regulon activity in each cell was calculated with the AUCell computational method (Aibar et al., 2017). By fitting regulon activity changes with an impulse model, we determined their temporal dynamics. As expected, the sequential changes of the regulon activity remarkably reflected the expression dynamics of the corresponding transcription regulators (Fig. 7G, 8A, 8B). The congruent changes of gene expression and regulatory state provide additional evidence for the important role of these transcription factors in the development of the rostral thalamus. We calculated the binary (on or off) activity of regulons and selected those that were enriched for each lineage (Fig. 8C and Fig. S5A-F). Furthermore, we generated a gene regulatory network (Fig. 8D). Therefore, combining URD and gene regulatory network analysis uncovered the transcriptional cascades that accompany the development of progenitors into differentiated cells and highlighted both previously characterized and newly identified trajectory-enriched genes.

**Figure 8.**
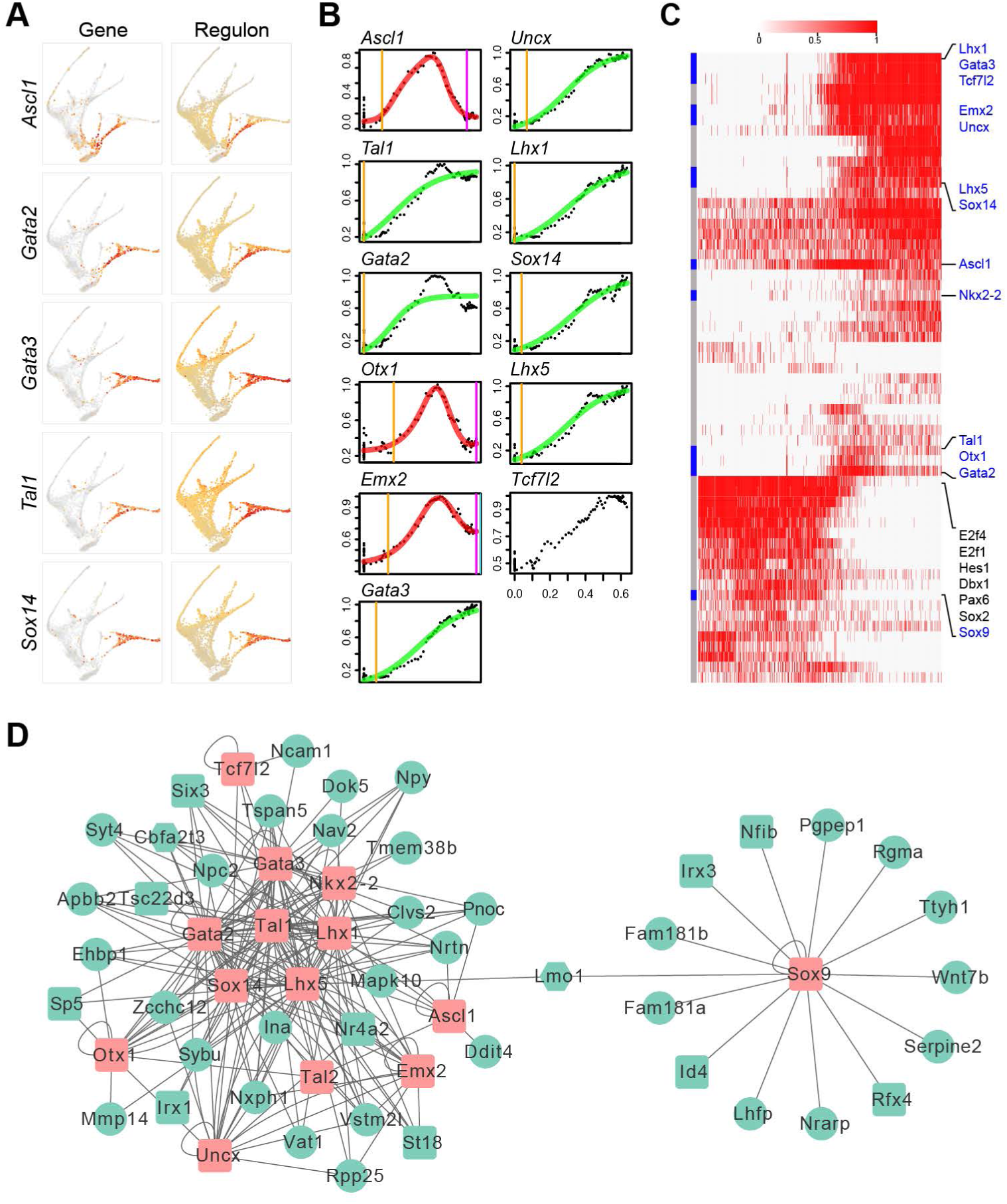
Gene regulatory networks underlying the caudal thalamic development. (**A**) Expression of genes (left) and regulons (right) overlaid on the trajectory tree. (**B**) Temporal changes of the regulon activity in association with the development of the rostral thalamus. (**C**) Heatmap showing the binary activity of regulons in the rostral thalamic cells ordered according to pseudotime (increasing from left to right). (**D**) Predicted network of the key regulators (pink) and their targets (green). Transcription factors are shown in rectangle.

In conclusion, our current study has characterized the molecular and cellular heterogeneity of the developing mouse diencephalon. We have identified key developmental regulators and their associated regulatory networks that define specific cell lineages. The resulting compendium of gene expression data provide opportunities to use genetic studies to interrogate the mechanisms that control cell fate decision and cell differentiation in the developing diencephalon. In addition, our approach by combining multiple state-of-the art statistical tools provides a framework with which we can classify cell types and cell states, study spatial pattern of the gene expression, and reconstruct the specification trajectories in highly complex developmental system with scRNAseq data.

## MATERIALS AND METHODS

### Generation and maintenance of mouse lines

All animal procedures described herein were approved by the Animal Care Committee at the University of Connecticut Health Center. Generation of *Gbx2*^*creER/+*^ (*Gbx2*^*tm1*.*1*)*cre/ERT2)Jyhl*^/J, #022135) mice were described previously (Chen et al., 2009). PCR conditions and primer information are provided in http://www.jax.org. The mouse line was maintained on a CD1 genetic background (Charles River Lab, Wilmington, MA). Noon of the day when a vaginal plug was detected was designated as E0.5 in staging of embryos.

### Generation of single-cell suspensions

Brains of *Gbx2*^*creER/+*^ embryos were dissected in ice-cold phosphate buffered saline. Under a fluorescent dissecting microscope, the thalamus, which was labeled by enhanced green fluorescent protein (EGFP) expressed from the *Gbx2* locus, was dissected together with surrounding tissues. Tissues from four animals were pooled, cut into pieces smaller than 1 mm in dimension, and transferred to ice-cold MACS Tissue Storage Solution (Miltenyi Biotec, Somerville, MA). For cell dissociation, the storage solution was exchanged with RPMI 1640 Medium (Thermo-Fisher); tissue was pelleted and digested with 500 µl of pre-warmed Accumax (Innovative Cell Technologies) in a 1.5-ml tube at 37°C for 5-10 min. At the end of digestion, the tissue pieces were dissociated by gentle trituration with a wide-bore pipet tip. The cell suspension was added to a 100-µm cell strainer (Corning, Corning, NY), and it was collected and transferred to 1.5 ml of ice-cold resuspension buffer (Lebovitz L15 medium with 2% FBS, 25 mM HEPES, 2 mM EDTA). The cell clumps that were unable to pass through the filter were placed into a new 500 µl pre-warmed Accumax solution, and the digestion and filtering process was repeated twice. Following the dissociation, cells were stained with Trypan blue, counted and visualized with Countess II Automatic Cell Counter (ThermoFisher). The single cell suspension had over 77% viability.

### Single-cell RNA-sequencing library construction and sequencing

Libraries were prepared using the Chromium Single Cell Kit v1 (PN-120233, 10x Genomics) and sequenced on Illumina NextSeq 500. The raw reads were processed to molecule counts using the Cell Ranger pipeline (version 1.3.1, 10x Genomics) with default setting (Zheng et al., 2017).

### Cell clustering and classification

The raw UMI counts from Cell Ranger were processed with the Seurat R package (version 1.4.0.9) (Satija et al., 2015). Genes that were detected in less than three cells were removed. Cells in which?over?5% of the UMIs were mapped to the mitochondrial genes and those that contained over?2,500 genes were discarded. Library-size normalization was performed on the UMI-collapsed gene expression for each barcode by scaling the total number of transcripts per cell to 10,000. The data was then log2 transformed. In total, 7,365 cells and 14,387 genes (an average of 1,147 detected genes/cell) were used in cell clustering. On average, we obtained 68,275 sequencing reads per cells, which represented median 1,758 unique genes per cells. We followed the standard Seurat workflow as described in the Satiji lab (https://satijalab.org/seurat/) to perform cell clustering and marker gene detection. For direct partitioning of roof-plate cells, we used the new interactive plotting function of Seurat (version >2.0).

For cell cycle score assignment, Seurat’s *RunPCA* function was used to perform principal component analysis with a list of cell cycle dependent genes (Tirosh et al., 2016). The resulting first principal component, which correlated with the activity of known S and G2/M phase genes, was referred to as a ‘cell cycle score’. Cells with cell cycle score below −3 or above 3 were assigned as S or M phase, respectively.

### Examination of spatial structures of scRNAseq data

We created a new Seurat object by extracting clusters 13 and 15 from the full dataset and performed normalization, scaling and t-SNE analyses. As inputs to trendsceek, the position of the 4,261 cells in the t-SNE were used as location of the points, and the normalized expression matrix was used as marks of the points. We used 10,000 permutations of the marks to obtain the null distribution representing marks being conditionally independent of the spatial location of the cells. We selected significant genes with Benjamini–Hochberg-adjusted p ≤ 0.05 for at least one of the four statistic tests (Edsgärd et al., 2018).

### Inference of developmental protectories

We used Monocle 2 (Qiu et al., 2017) to perform pseudotime analysis. For the caudal thalamus, we generated a new Seurat object by extracting clusters 3, 6, 10, and 15. For the rostral pretectum, we created a new Seurat object by extracting clusters 5, 7, 9, and 16. The resulting Seurat objects were exported to Monocle. We reconstructed the developmental trajectories in these two lineages by following the standard procedure described in the Trapnell lab (http://cole-trapnell-lab.github.io/monocle-release/tutorials/). Temporal gene expression profiles were plotted using the ggplot2 package (Wickham, 2016).

To perform URD analysis, we followed the procedure described previously (Farrell et al., 2018). We assigned presumptive apical progenitors (mostly clusters 15 and 16) as the root, and postmitotic neurons (clusters 1-5 and 7) as tips, resulting a branching trajectory tree. The URD *aucprTestAlongTree* function was used to identify the “cascade genes” that were progressively increased along each trajectory. To identify lineage-specific genes, we used all the cascade genes for each lineage and selected those that were highly enriched in the given lineage compared to the rest of the diencephalon. We used the area under the precision-recall curve for this gene (AUC) was greater than the area expected for a random feature by a factor of 3. To examine the enrichment of lineage markers in *Gbx2* target genes, we performed hypergeometric tests using the R package piano (Varemo et al., 2013).

For the genes identified as members of a gene expression cascade, we used the URD *geneCascadeProcess* function to smooth their expression using a moving window average (18 cells per window, moving 5 cells at a time) along the trajectory. The value was then scaled to the maximum window average for each gene in the trajectory. The smoothed expression of each gene was fit with an impulse model to determine its temporal expression dynamics (Farrell et al., 2018).

### Inference of single-cell gene regulatory network

We used the SCENIC pipeline (Aibar et al., 2017) to determine the gene regulatory network in each diencephalic lineage. We first applied GENIE3 (Huynh-Thu et al., 2010) to gene count matrix to construct a co-expression network. As input to GENIE3, we filtered genes with at least 3 counts in 10% cells, and cells with at least 90.15 UMI, resulting a final matrix of 8,320 genes and 7,162 cells. We used the *RcisTarget* algorithm to select potential direct-binding targets (regions) based on DNA-motif analysis, and then the *AUCell* algorithm to calculate the network activity in each individual cell, resulting in an activity matrix in which the columns represent cells, and the rows regions (Aibar et al., 2017). Using a default threshold, we converted the regulon activity to on/off binary activity matrix, which was then used to produce heatmaps. We examined the temporal changes of regular activity by applying the URD’s *geneCascadeProcess* function to the continuous AUC matrix. The gene regulatory network was exported to and modified in Cytoscape (version 3.7.0) (Shannon et al., 2003).

### Data availability

The sequencing files and raw gene count matrix have been deposited in NCBI’s Gene Expression Omnibus and are accessible through accession number GSE122012. All the computer codes and necessary data associated with the manuscript are provided in the supplementary information, and are also available at https://github.com/JLiLab/scRNAseq_Diencephalon.

## Supporting information

## ACKNOWLEDGEMENTS

We are grateful to the JAX-UConn Single Cell Genomics Center. We thank Drs. Paul Robson and Mohan Bolisetty for their advice and technical assistance in scRNAseq experiments. We also thank Mr. Elliott Wilion for language editing and proofreading.

## SUPPLEMENTARY DATA

**Figure S1.**
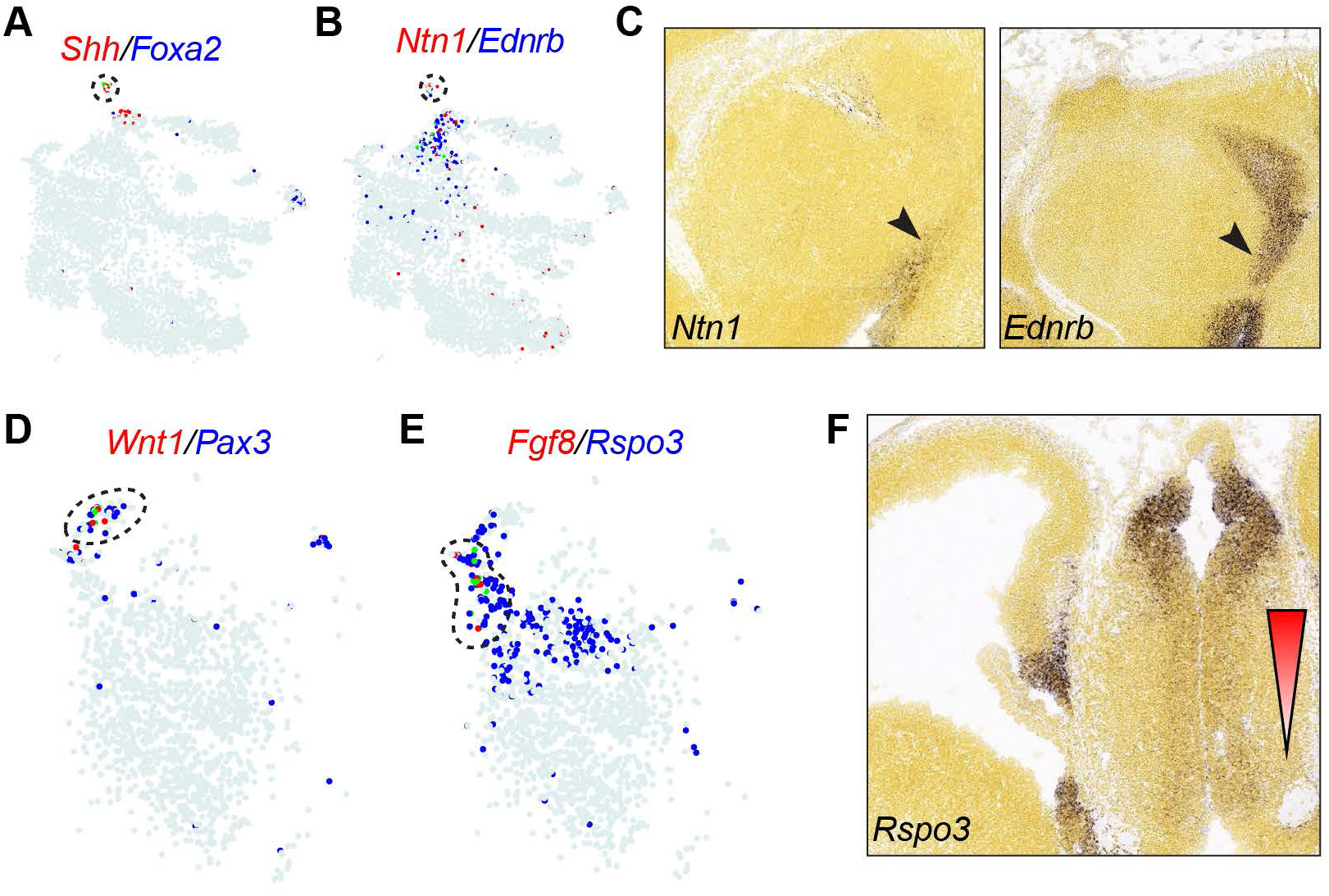
Assignments of cell groups corresponding to the basal plate and the roof plate. (**A** and **B**) t-SNE plotting of the expression of markers for cluster 12 (circled by dashed lines). (**C**) *In situ* hybridization on sagittal sections of E13.5 mouse brain. (**D** and **E**) t-SNE showing expression of *Fgf8* and *Wnt1* in two distinct subgroups of cells (circled by dashed lines) within cluster 13. (**F**) *In situ* hybridization on coronal sections of E13.5 mouse brain. The triangle indicates the decreasing gradient of *Rspo3* expression in the dorsal-to-ventral direction. Images in C and F are from Allen Developing Mouse Brain Atlas.

**Figure S2.**
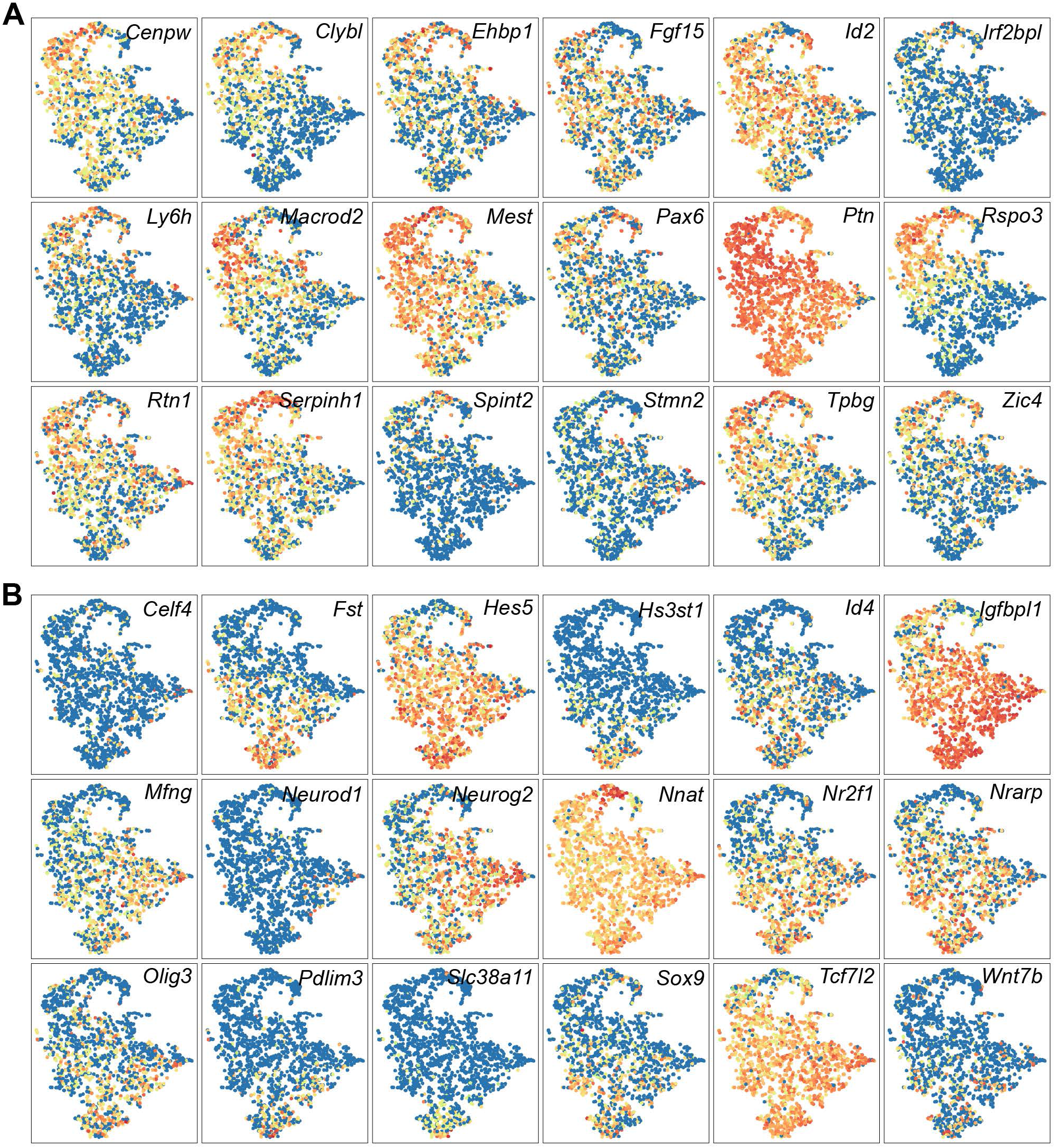
Trendsceek analysis of spatial expression patterns with scRNAseq data. (**A** and **B**) Trendsceek-identified significant genes that exhibit dorsal^high^-ventral^low^ (A) or dorsal^low^-ventral^high^ (B) gradients in the p2 ventricular zone.

**Figure S3.**
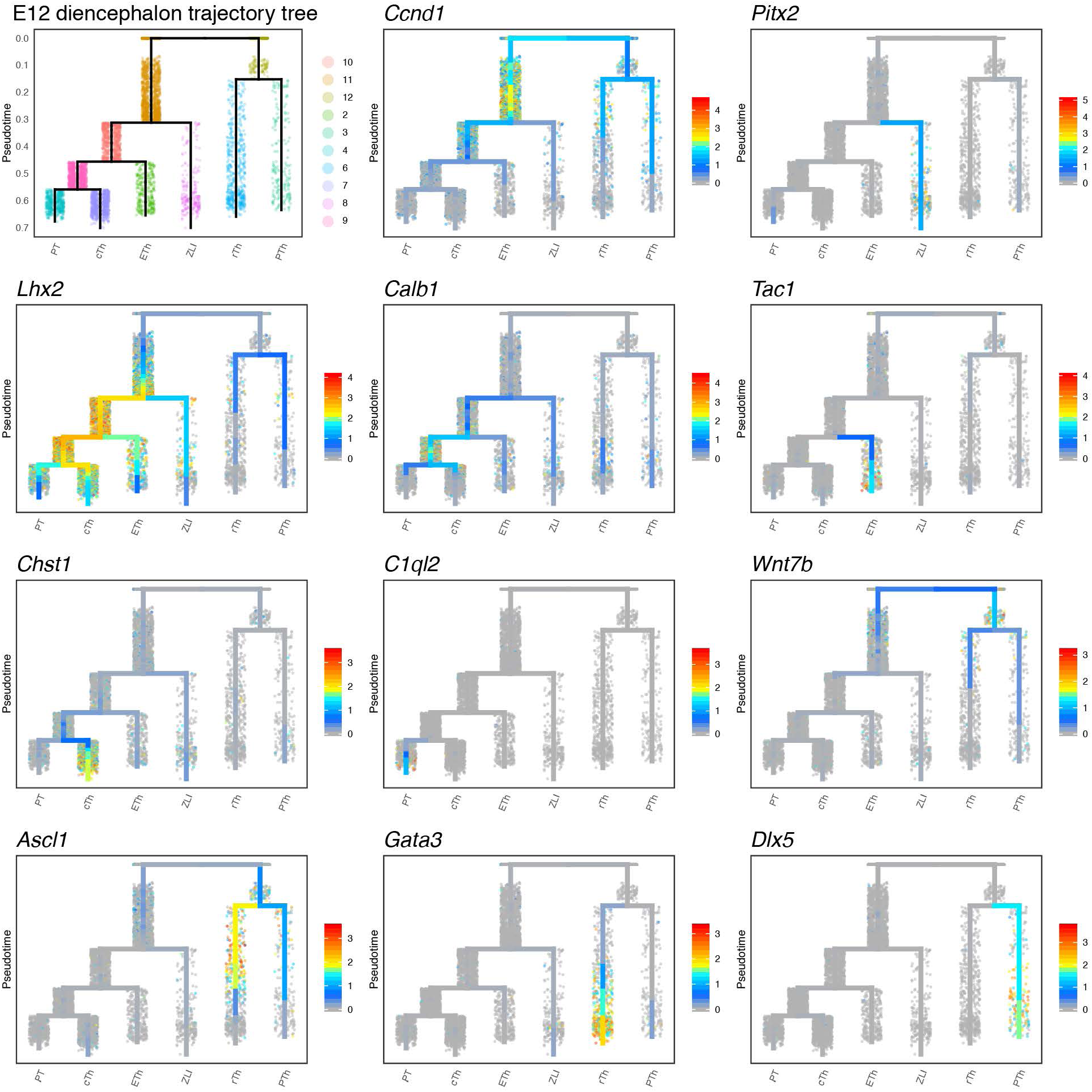
Expression of marker genes in the URD-inferred trajectory tree. Variable expression of markers highlighting distinct segment(s) of the trajectory as expected.

**Figure S4.**
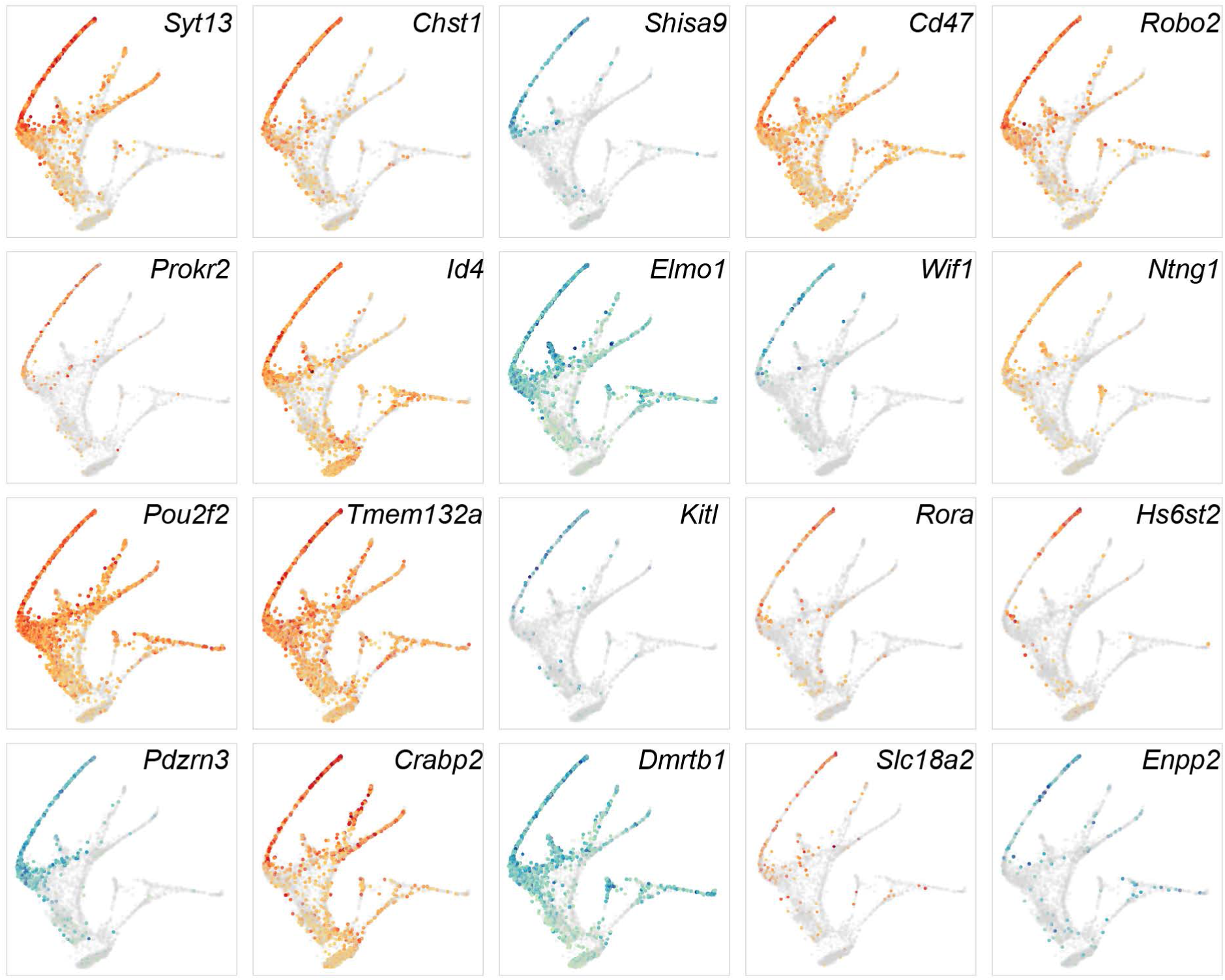
Expression of thalamus-specific genes overlaid on the trajectory tree. Note the gradual increase and the enrichment of gene expression along the trajectory corresponding to the caudal thalamus. Genes that are previously identified as thalamus-specific genes (Mallika et al., 2015; Suzuki-Hirano et al., 2011) are plotted in orange, whereas new markers are plotted in blue/green.

**Figure S5.**
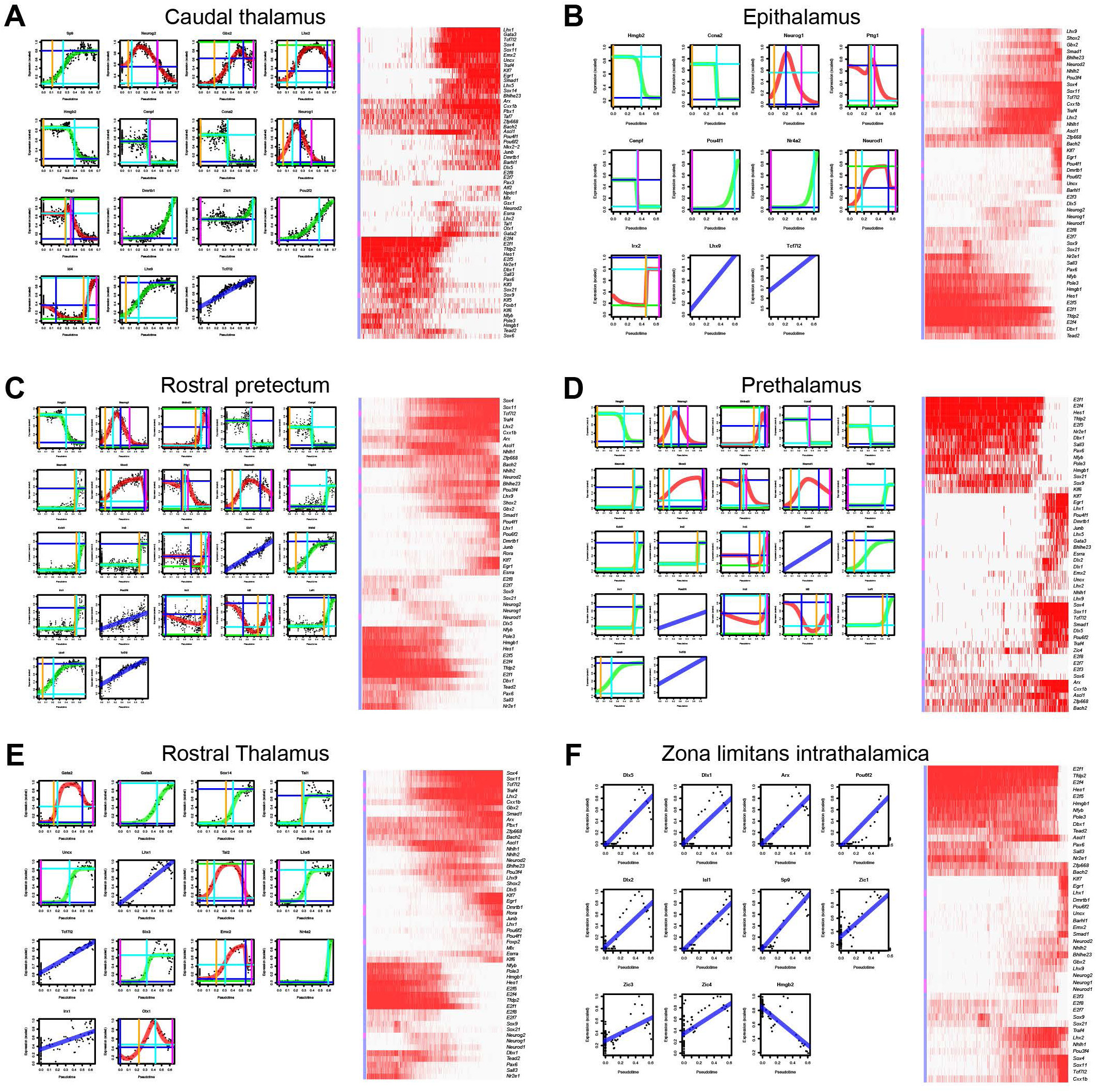
Temporal dynamics of key transcription factors and the associated regulons in different diencephalic lineages. **(A-F)** Temporal expression of transcription factors (left), and binary regulon activities (right) in six lineages of the diencephalon. In the x axis, cells are arranged according to pseudotime (increases from left to right). Each dot represents the moving-window average of expression levels; lines show impulse response fitting: blue for genes that change at a constant rate, green for a single sigmoid impulse, and red for a convex or concave impulse; vertical lines indicate the onset (orange) and offset (magenta) of regulon activation. The transcription factors that are significantly changed during lineage specification are indicated (in red) by the color bar on the left side of the heatmap.

Table S1. Excel file of mark genes of each cluster. (**A**) Differentially expressed genes identified by Seurat (first spreadsheet). (**B**) Top 20 markers for each cluster (second spreadsheet). (**C**) Marker genes for neural progenitor cells (NPC), neuronal precursors (NPre), and postmitotic neurons (N) (third spreadsheet). Column A: gene symbol; B: transcription factor; column C: description of gene name; column D: unadjusted p value; Column E: adjusted p value based on bonferroni correction using all genes in the dataset; Column F: log2 fold-change of the average expression between the two groups (positive values indicate that the gene is more highly expressed in the first group); Column G: The percentage of cells where the gene is detected in the testing cluster; Column H: the percentage of cells where the gene is detected in the rest of the dataset; Column I: difference of the percentage of cells where the gene is detected; Column J: cluster identification; Column K: annotation of the cell cluster.

Table S2. Excel table of cascade genes of six diencephalic lineages. (A) Cascade genes identified by URD’s *aucprTestAlongTree* function (first spreadsheet). (B) Lineage specific markers (second spreadsheet)

